# Identification of a host collagen inducing factor from the excretory secretory proteins of *Trichinella spiralis* using immunoscreening

**DOI:** 10.1101/319483

**Authors:** Mi Kyung Park, Hae-Jin Kim, Min Kyoung Cho, Shin Ae Kang, So Young Park, Se Bok Jang, Hak Sun Yu

**Affiliations:** Department of Parasitology School of Medicine, Pusan National University, Yangsan, 50612, Republic of Korea; Department of Molecular Biology, College of Natural Sciences, Pusan National University, Busan 46241, Republic of Korea

**Keywords:** *Trichinella spiralis*, serine protease, collagen inducing factor, immunoscreening

## Abstract

**Background:** In a previous study, we found that *Trichinella spiralis* excretory and secretory proteins (ES-P) most likely activate collagen synthesis via TGF-β/Smad signaling, and this event could influence collagen capsule formation.

**Methodology/Principal Findings:** In order to identify the specific collagen inducing factor, ES-P was fractionated by a Superdex 200 10/300 GL column. We obtained three large fractions, F1, F2, and F3, but only F3 had collagen gene inducing ability. After immunoscreening, 10 collagen inducing factor candidates were identified. Among them, TS 15-1 and TS 15-2 were identical to the putative trypsin of *T. spiralis*. The deduced TS 15-1 (M.W. = 72 kDa) had two conserved catalytic motifs, an N-terminal Tryp_SPc domain (TS 15- 1n) and a C-terminal Tryp_SPc domain (TS 15-1c). To determine their collagen inducing ability, recombinant proteins (rTS 15-1n and rTS 15-1c) were produced using the pET-28a expression system. TS 15-1 is highly expressed during the muscle larval stage and has strong antigenicity. We determined that rTS 15-1c could elevate collagen I via activation of the TGF-β1 signaling pathway *in vitro* and *in vivo*.

**Conclusion/Significance:** In conclusion, we identified a host collagen inducing factor from *T. spiralis* ES-P using immunoscreening and demonstrated its molecular characteristics and functions.

**Author Summary:** *Trichinella spiralis* can make collagen capsules in host muscle cells during its life cycle, which encapsulates muscle stage larvae. Many investigators have tried to reveal the complex mechanism behind this collagen capsule architecture, and it has been suggested that several serine proteases in excretory-secretory proteins of the parasite are potential collagen capsule inducing factors. In addition, collagen synthesis is activated through the TGF-β/Smad signaling pathway and these events are closely related with protease activated receptor 2 which was activated by various serine proteases. In this study, we isolated and characterized a collagen gene expression inducer from *T. spiralis* ES-P using immunoscreening and investigated the candidate protein for its usefulness as a wound healing therapeutic agent.

## Introduction

*Trichinella spiralis* can make collagen capsules (nurse cell formation) in host muscles during their life cycle that surround muscle stage larvae and might protect the larvae from the host immune system. This phenomenon can be understood as the parasite creating a simple structure to protect itself, but when examined closely, numerous different mechanisms are involved in this stage of the parasite’s life. Division of the host muscle cell nucleus, regulation of host cell cycling, huge elevation of host collagen gene expression, and generation of new blood vessels around the collagen capsule are observed during nurse cell formation by *T. spiralis* [1-4]. The process of nurse cell formation induces de-differentiation, cell cycle re-entry, arrest of infected muscle cells, and activation, proliferation, and differentiation of satellite cells. These events are very similar to those occurring during muscle cell regeneration and repair [2].

Tissue regeneration and repair are of great interest in the research field of wrinkle improvement of aging skin tissue. Skin collagen, in the form of elongated fibrils, provides skin strength and elasticity. The gradual loss of collagen in skin with aging results in wrinkles and other signs of skin aging. The content of type I collagen, the major collagen in the skin and a marker of collagen synthesis, is decreased by 68% in old skin versus young skin, and cultured young fibroblasts synthesize more type I collagen than old cells [5]. The main reason for the decreased production of collagen was found to be due to decreased synthesis of type I collagen mRNA, and there was a three-fold reduction in the steady-state level of type I collagen mRNA in senescent fibroblasts [6]. Another fundamental mechanism for age-related decrease in collagen synthesis is the TGF-β-induced signaling pathway of collagen synthesis. TGF-β is the most potent direct stimulator of collagen production. Moreover, TGF-β is central to the process of wound healing and fibrosis formation [7]. It is well understood that Smad signaling pathways are activated downstream of TGF-β via receptor-mediated Smad phosphorylation, and these pathways activate collagen synthesis. Thus, TGF-β plays an important role in regulating extracellular matrix synthesis [8].

In a previous study, we found that *T. spiralis* excretory and secretory proteins (ES-P) most likely activate collagen synthesis via TGF-β/Smad signaling, and this event could influence collagen capsule formation [9]. These events were closely related with protease activated receptor 2 (PAR2), which was activated by various serine proteases [9]. However, the question of which protease in *T. spiralis* ES-P has a role of collagen gene expression of host muscle cells is still unanswered. The identification of a specific collagen gene inducer from *T. spiralis* could be exploited as a therapeutic and/or cosmetic agent.

In this study, we isolated and characterized the collagen gene expression inducer from *T. spiralis* ES-P by immunoscreening and investigated the candidate for its usefulness as a wound healing therapeutic agent.

## Materials and Methods

### Extraction of ES-P from muscle larvae

The *T. spiralis* strain (isolate code ISS623) used in this study has been maintained in our laboratory via serial passage in rats. Muscle larvae were isolated from *T. spiralis* infected mice (4 weeks after infection) and ES-P from cultured muscle larvae was obtained according to the previously reported method [9].

### Mouse experiments

Twenty female C57BL/6 mice at the age of 6 weeks and twenty female 14 week-old mice were purchased from Samtako Co. (Gyeonggi-do, Korea). The skin of the left ear of each mouse was treated with *T. spiralis* ES-P (30 μg) in PBS every day for 14 days, and that of the right ear was treated with PBS. The mice were housed in a specific pathogen-free facility at the Institute for Laboratory Animals of Pusan National University.

### Fractionation of ES-P

ES-P (5 mg) in 10 ml PBS was applied to a Superdex 200 10/300 GL column (GE Healthcare, Uppsala, Sweden). The flow rate was 0.25 ml/min. Each 0.5 ml fraction was collected and protein quantity was measured by UV detection at 260 nm. Three big fractions, F1, F2, and F3, were acquired and used for collagen gene inducing experiments (Fig. 3A).

**Fig. 1.**
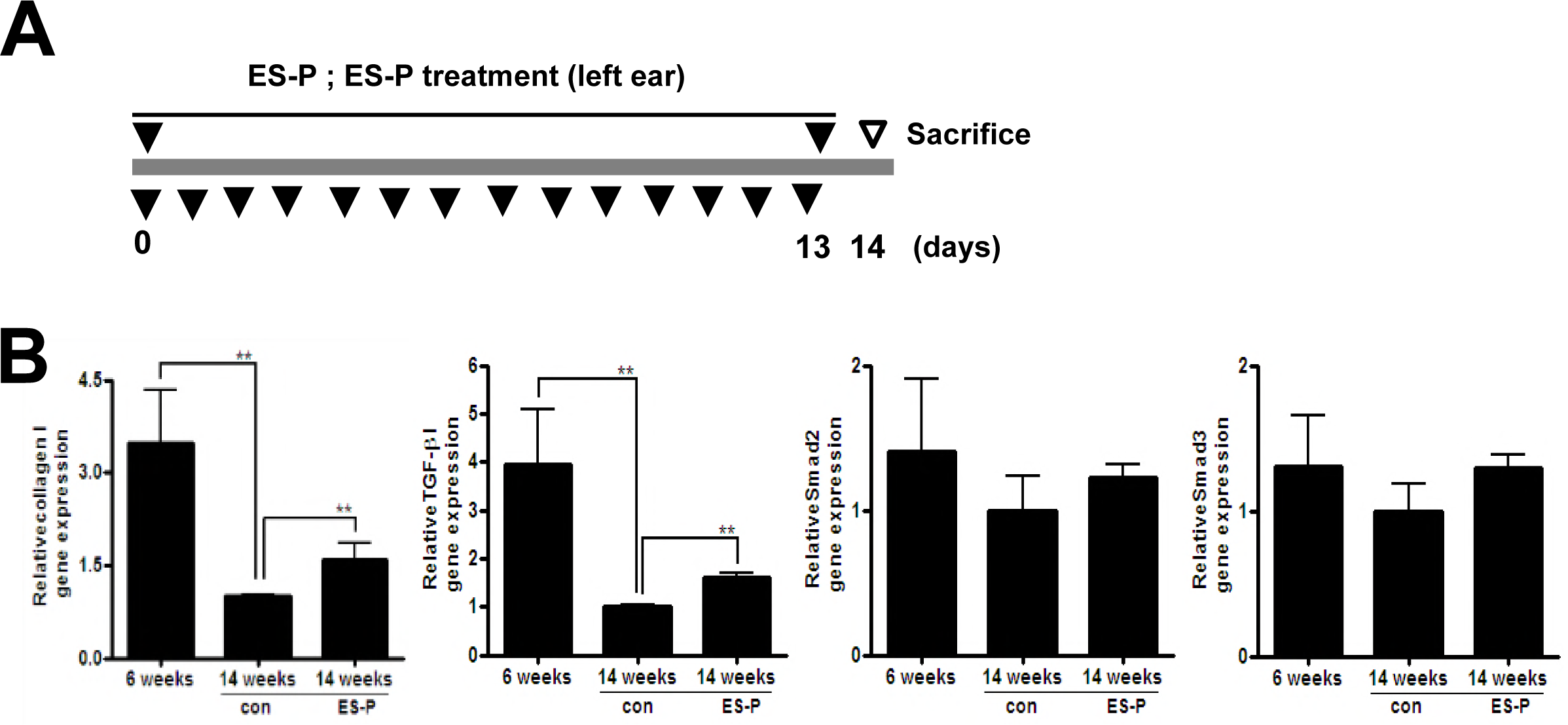
Expression levels of type I collagen, Smad2/3, and TGF-β1 genes after ES-P treatment in aged mice. The left ears of fourteen-week old mice were treated daily with *T. spiralis* ES-P (30 μg) for 14 days (A). After 14 days, left ears and non-treatment right ears were collected. Total RNA was extracted from each ear tissue and cDNA was constructed. The gene expression levels of *collagen I*, *TGF-β1*, and *Smad2/3* were analyzed by real-time PCR (B). (**; *P* < 0.01, n = 5 mice/group, these were representative results from three independent experiments).

**Fig. 2.**
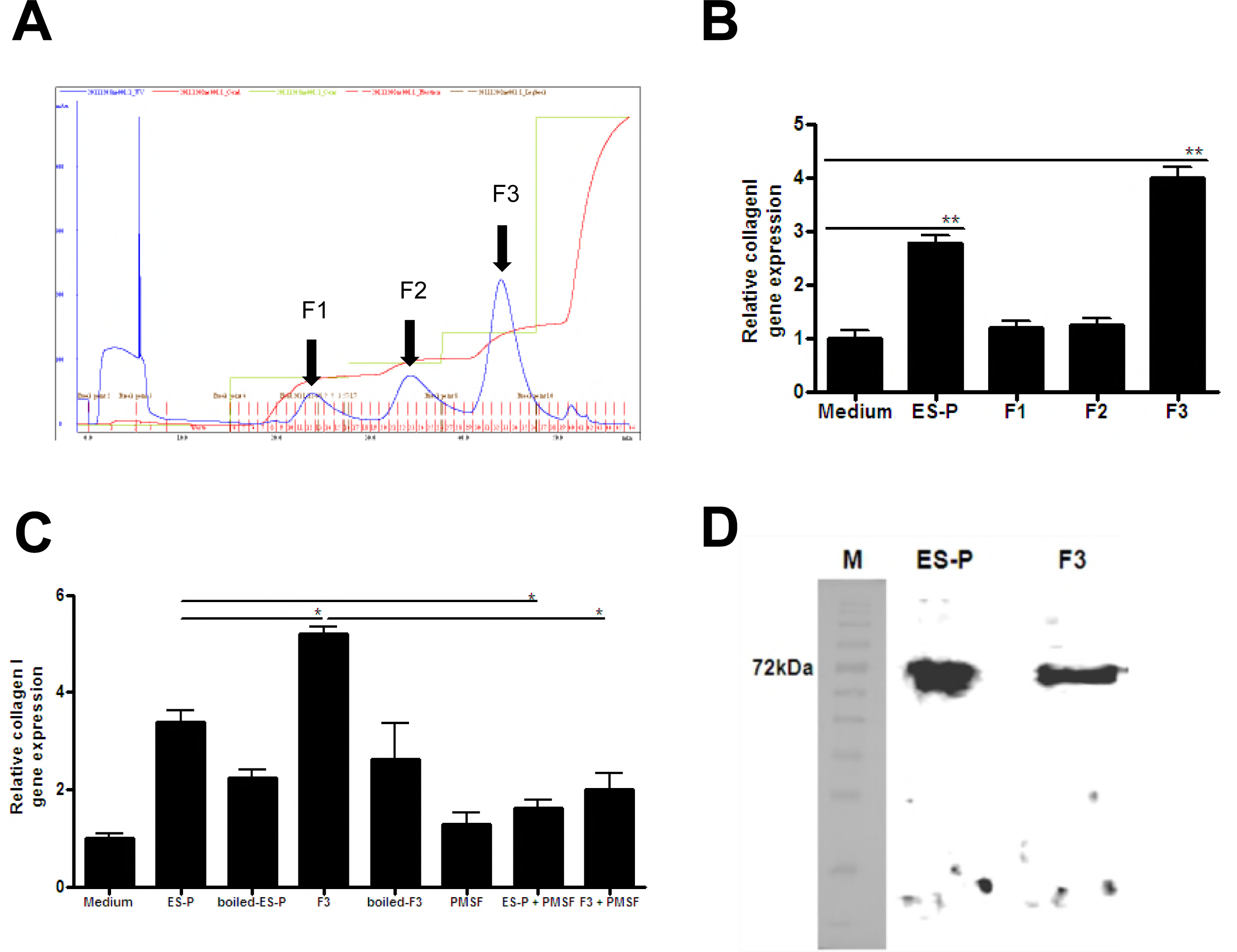
Fractionation of the ES-P by chromatography and type I collagen expression analysis. Chromatograhic profile of ES-P conducted by ion exchange fractionation (A). Type I collagen gene expression levels were analyzed after ES-P fractionation treatment (B). After MEF cell treatment with fraction 1, 2, and 3 (each at 1 μg/ml) for 2 hrs, total RNA was extracted from each sample and cDNA was synthesized. The gene expression levels of *collagen I* was analyzed by real-time PCR. Medium; cell culture medium, ES-P; *T. spiralis* ES proteins, F1; F1 fraction, F2; F2 fraction, F3; F3 fraction. Evaluation of collagen gene expression was closely related with protease activity (C). After treatment with the F3 fraction (1 μg/ml) in presence or absence of a protease inhibitor, or treatment of the boiled F3 fraction for 2 hrs, total RNA was extracted from each MEF cell sample and cDNA was constructed. Boiled ES-P; treatment with *T. spiralis* ES protein boiled for 10 min at 90°C, boiled-F3; treatment with F3 boiled for 10 min at 90°C, PMSF; only PMSF treatment (final conc. 1 mM), ES-P+PMSF; *T. spiralis* ES proteins and PMSF treatment, F3+PMSF; F3 fraction and PMSF treatment. Western blot analysis of ES-P and F3 with polyclonal α-F3 antibody. Ten micrograms of ES-P and F3 protein samples were loaded on an SDS gel and western blot analysis was performed. The α-Ts F3 antibody (1:500 dilution) was used as primary antibody. The α-rat IgG-HRP conjugate was used as the secondary antibody at a 1:5000 dilution. (*; *P* < 0.05, **; *P* < 0.01, n = 5 mice/group, these were representative results from three independent experiments).

**Fig. 3.**
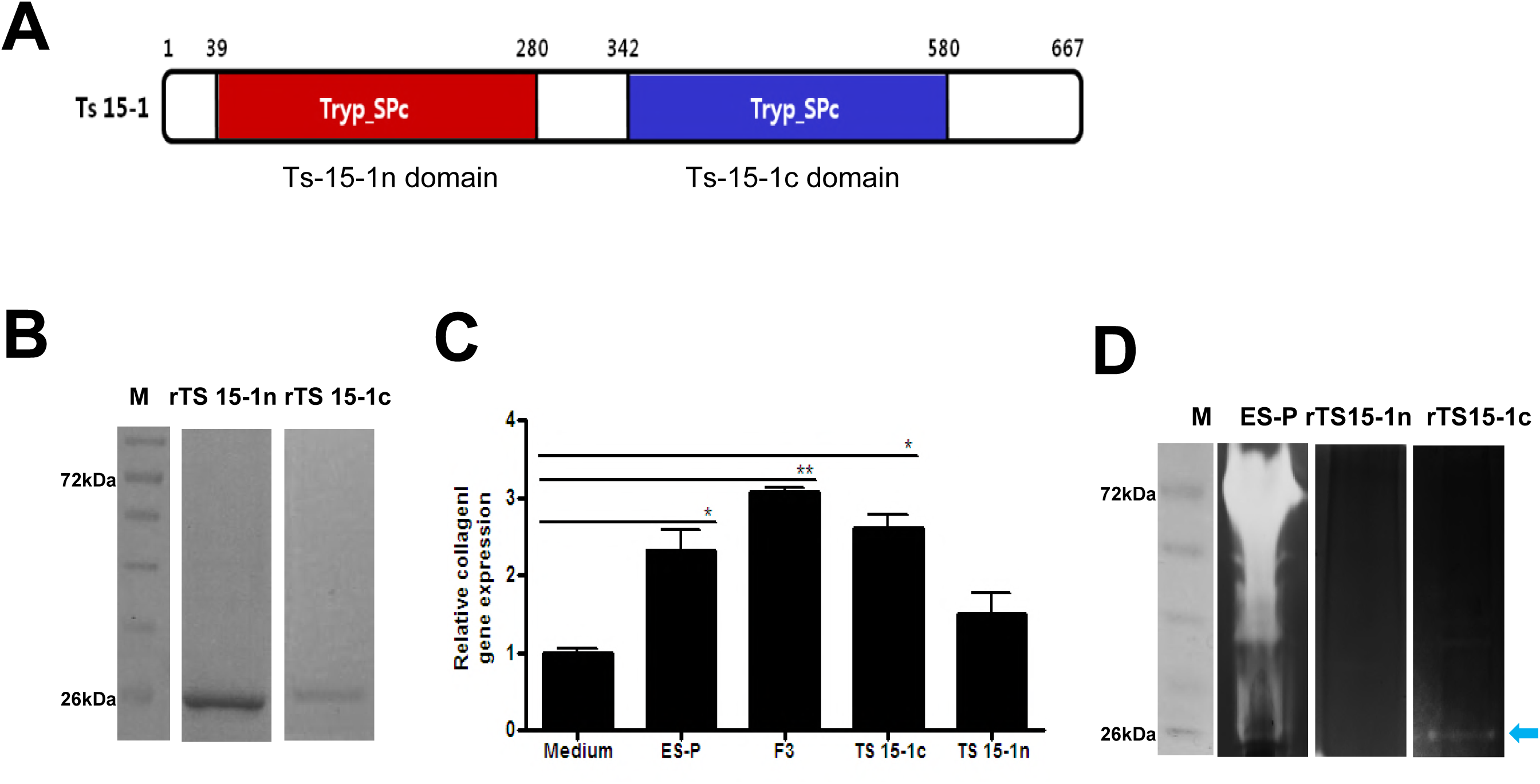
Molecular structure and characterization of TS 15-1. Schematic diagram showing the domains of the full-length TS 15-1. TS 15-1 consists of two trypsin domains (A). (red; N-terminal Tryp_SPc domain, blue; C-terminal Tryp_SPc domain). SDS-PAGE analysis of recombinant proteins rTS 15-1n and rTS 15-1c (B). Comparison of collagen gene inducing abilities of the recombinant proteins and ES-P (C). After MEF cell treatment with ES-P, F3, rTS 15-1n, rTS 15-1c (each at 1 μg/ml) for 2 hrs, expression levels of *collagen I* were analyzed by real-time PCR. Proteolytic activity evaluation of two recombinant proteins (D). One microgram each of ES-P, rTS 15-1n, and rTS 15-1c was used for gelatin-zymography to determine protease activity (D). Arrow indicates protease activity (*; *P* < 0.05, **; *P* < 0.01, n = 5 mice/group, these were representative results from three independent experiments).

### Cell culture and *in vitro* stimulation

Mouse embryonic fibroblast (MEF) cells were isolated from C57BL/6 mouse fetuses 10 days after fertilization. MEF cells were incubated in DMEM (Difco) with 5% FBS and 5 × 10^5^ cells were plated in 24-well plates and incubated overnight at 37°C in 5% CO_2_. Cells were treated with samples (final conc. 1 μg/ml) with or without PMSF (serine protease inhibitor, Sigma-Aldrich, USA).

### Real-time PCR

Homogenized ear tissues were mixed with TRIzol (Invitrogen, Germany), and RNA extraction and cDNA synthesis (Invitrogen, Germany) was performed in accordance with the manufacturer’s protocols. Expression levels of several genes were determined with real-time RT-PCR using the iCycler^TM^ (Bio-Rad laboratories Inc., USA) real-time PCR machine. Primer sequences for *collagen I, TGF-β, smad2, smad3*, and *GAPDH*, and PCR conditions were identical to those mentioned in the previous study 9]. The primer sequences for the putative trypsin (TS 15-1c) were 5′- TTG GAA TGA CGC TGA TTG -3′, 5′- GTG GCT TAT GAT GGT AGG AGA AT -3′.

### Production of polyclonal antisera for F3 (α-F3 antibody)

Female four-week-old Wistar rats were purchased from Samtako Co. (Gyeonggi-do, Korea). Rats were immunized subcutaneously with a combination of 250 μg F3 fractions (in 0.5 ml PBS) and 0.5 ml Freund’s adjuvant (Sigma-Aldrich, USA) at 0 and 2 weeks (booster – 2^nd^ subcutaneous injection). One week after their final booster, rats were sacrificed and serum was obtained.

### Immunoscreening of cDNA library

A cDNA library generated from 60,000 plaques forming units of *T. spiralis* muscle larvae was screened with the α-TS F3 antibody. Immunoscreening was performed in accordance with the manufacturer’s protocols (Stratagene, USA). Briefly, after primary and secondary screening, positive plaques were picked and the phagemids were prepared by *in vivo* excision. The phagemids were transformed into XL1-Blue MRF cells. Clones were selected based on blue-white color selection of the colonies grown on LB-ampicillin agar plates. The plasmid harboring the cDNA inserts were then extracted using a plasmid DNA purification system (Cosmogenetech, Seoul, Korea). The cDNA inserts were then sequenced using the primer for T3 promotor (Cosmogenetech, DNA sequencing service, Seoul, Korea) and compared against the GenBank database.

### Expression of the TS 15-1c domain and TS 15-1n domain

The *TS 15-1c* (C-terminal serine protease domain) and *TS 15-1n* (N-terminal serine protease domain) genes were amplified and ligated into the pET-28a expression vector. After gene ligation, the constructed plasmids were transformed into *Escherichia coli* BL21. The expressed TS 15-1c and TS 15-1n proteins were loaded on a 10% SDS-PAGE gel.

### Multiple sequence alignment and phylogenetic analysis

Multiple sequence alignments were performed with GeneDoc 5.0 and phylogenetic analysis was conducted in Clustal Omega. The optimal tree with the sum of branch length = 0.1 is shown. The tree is drawn to scale, with branch lengths in the same units as those of the evolutionary distances used to infer the phylogenetic tree. The evolutionary distances were computed using the Poisson correction method and the units were the number of amino acid substitutions per site.

### Western blotting

Ten micrograms of the ES-P and F3 fractions were loaded into each well of a 10% acrylamide SDS-PAGE gel and the proteins were separated at 100 V for 90 min. The loaded proteins were transferred onto a nitrocellulose membrane (Amersham Biosciences, Little Chalfont, UK) and blocked with 5% skim milk in TBST at 4°C overnight. Then, the membrane was incubated with primary antibody (α-F3 antibody, rat serum, 1:500) in 5% skim milk in TBST for 2 hrs at room temperature. The secondary antibody, α-rat IgG-HRP conjugate (Sigma, Seoul, Korea) was used at 1:5000 dilution for 1 hr at room temperature. HRP was detected using an ECL substrate (Amersham Biosciences, Uppsala, Sweden).

### Immunohistochemistry

Paraffin-embedded *T. spiralis* muscle stage larvae were de-paraffinized and hydrated. For antigen retrieval, slides were immersed in citrate buffer (0.01 M, pH 6.0) and heated twice in a microwave (700 W or ‘high’) for 5 min. Slides were then quenched with endogenous peroxidase by incubation in a 3% hydrogen peroxide solution for 5 min and were washed three times in PBS for 5 min each. Slides were immuno-stained with primary antibody (α-TS 15-1c antibody that was produced according to the polyclonal antisera method; 1:500 dilution) at 4°C overnight. After primary antibody incubation, slides were washed three times in PBS for 5 min each and were incubated with secondary antibody for 1 hr at room temperature. Slides were subsequently washed four times in PBS for 5 min each and the color reaction was developed with Dako’s EnVisionTM System (DAKO, Carpinteria, CA, USA). Slides were stained with 3,3'-diaminobenzidine (DAB) and counterstained with Meyer’s hematoxylin (DAKO, Carpinteria, CA, USA) for 20 sec, dehydrated, and mounted with Permount (Fisher Scientific, Pittsburgh, PA, USA). Background staining was checked by the immuno-staining of all slides without primary antibody and antibody specificity was confirmed with positive controls.

### Statistical analysis

All experiments were performed three times for confirmation of statistical significance. Mean ± standard deviation (SD) was calculated from data collected from individual mice. Significant differences were determined using one-way or two-way analysis of variance. Statistical analysis was performed with GraphPad Prism 5.0 software (GraphPad Software Inc., CA, USA).

### Ethics statement

The study was performed with approval from the Pusan National University Animal Care and Use Committee (IACUC protocol approval; PNU-2016-1175), in compliance with ‘‘The Act for the Care and Use of Laboratory Animals’’ of the Ministry of Food and Drug Safety, Korea. All animal procedures were conducted in a specific pathogen-free facility at the Institute for Laboratory Animals of Pusan National University.

## Results

### Type I collagen gene expression in an aging mouse model

In order to understand the collagen gene inducing effect of ES-P, transcription and protein expression levels of type I collagen and TGF-β1 signaling related proteins were compared in ear tissues of 6 and 14 week-old mice that had or had not received ES-P treatment (Fig. 1A). The expression levels of the *collagen I* and *TGF-β1* genes of the 14 week-old mice were significantly decreased compared to those of the 6 weeks mice. However, those of the ES-P treated 14 week-old mouse group were significantly increased compared to un-treated 14 week-old mice (Fig. 1B).

### Fractionation of ES-P and evaluation of collagen gene induction of major fractions

To identify type I collagen inducing factors from the *T. spiralis* ES-P, the ES-P was fractionated to several fractions including three big fractions by gel chromatography (Fig. 2A). The three major fractions obtained were named F1 (about 140 kDa – 100 kDa), F2 (about 120 kDa - 80 kDa), and F3 (about 90 kDa - 50 kDa) respectively (Fig. 2A). In order to determine which major fraction had collagen gene expression inducing factors, each fraction was used to treat MEF cells, and expression levels of the type I collagen gene were measured. After F1, F2, and F3 treatment, only F3 treated MEF cells had significantly increased type I collagen gene expression (Fig. 2B). Moreover, expression levels of *collagen I* in F3 treated MEFs was higher those of ES-P treated MEFs.

### Protein activity evaluation and size determine of F3

In a previous study, the collagen inducing ability of ES-P was closely related with serine protease activity. In order to determine whether the collagen inducing ability of F3 is related with serine protease activity, we evaluated the collagen inducing ability of F3 following pre-treatment with a serine protease inhibitor, PMSF, on MEF cells. *collagen I* and *TGF-βI* gene expression levels were significantly decreased in MEF cells pre-treated with PMSF, the serine protease inhibitor. In addition, the expression levels were also not increased when treated with boiled F3 (Fig. 2C). In order to confirm of existence and expression levels of F3 in ES-P, α-F3 polyclonal antibody was used against ES-P and F3 in a western blot analysis. The presence of a strong band was observed at 60 - 80 kDa in both the ES-P and F3 samples (Fig. 2D).

### Immunoscreening of a cDNA library from *T. spiralis* muscle larva by α-F3 antibody

In order to identify the collagen gene inducing factors from *T. spiralis* ES-P, immunoscreening was conducted against the *T. spiralis* muscle larvae Express cDNA library with the α-F3 antibody. Thirty-five positive plaques were detected in primary screening, among them, 10 plaques were confirmed by second screening (Supplementary Fig. 1). These plaques were amplified and processed in an *in vivo* excision step. All the insert DNA from the 10 positive clones were sequenced and their amino acid sequences were determined. Two insert DNA fragments (TS 15-1 and TS 15-2) were similar to a putative trypsin of *T. spiralis* with 90% identity. Another clone (TS 15-3) was matched to a nuclear receptor-binding protein of *T. spiralis* with 35% of identity. Another clone (TS-16-1) was matched to a putative BTB/POZ domain protein of *T. spiralis* with 63% identity. The remaining 6 insert clones were not matched with any previously known genes.

### Molecular characterization of *TS 15-1*

Collagen inducing factors in ES-P and F3 had serine protease activity and were measured to be about 60 - 72 kDa in size. After evaluation of the size and serine protease activity of positive clone matched genes, the *TS 15-1* gene was selected for downstream identification of the collagen inducing factor. The *TS 15-1* fragment was 2,004 bp long and encoded a 667 amino acid protein, and the molecular weight and pI was calculated as 71.6 kDa and 8.83. The deduced TS 15-1 protein has two conserved catalytic motifs, an N-terminal Tryp_SPc domain (TS 15-1n) and a C-terminal Tryp_SPc domain (TS 15-1c) (Fig. 3A). The TS 15-1n and TS 15-1c peptides were composed of 238 amino acids and 239 amino acids respectively, and the molecular weight was calculated to be 26.1 kDa and 26.2 kDa respectively. Unfortunately, collection of recombinant full length of TS-15 protein was very difficult because it was produced in very small amounts in this system. We conducted recombinant protein expression of the N and C terminal domains (about 26 kDa, Fig. 3B) and evaluated their collagen gene inducing ability. *collagen I* expression levels in the recombinant TS 15-1c protein treated cells were significantly increased compared to a media control. However, recombinant TS 15-1n protein treated cells were not significantly changed compared to those of medium treated cells (Fig. 3C). In order to know the protease activity of both recombinant proteins, zymogram analysis was conducted. A collagen digested clear zone was detected around ~26 kDa in the recombinant TS 15-1c protein lane, but the clear zone was not detected with the recombinant TS 15-1n protein (Fig 3D).

### Molecular structure of TS 15-1c

In this study, we determined a molecular model by homology modeling based on the structure of another serine protease (PDB ID: 1KYN, 1-235). In TS 15-1c (G342 - T580), predictions of the active sites (H389, D444, and S533) and the substrate binding sites (G527, S553, and G555) are shown in blue and red letters, respectively (Fig. 4A). Interestingly, these results indicated that these sites interact with inhibitors and ligands. Most residues in these regions had negative charges in a globular fold (Fig. 4B). In addition, the results of phylogenetic analysis indicated that TS 15-1c was closely related with the granzyme B category of serine proteases.

**Fig. 4.**
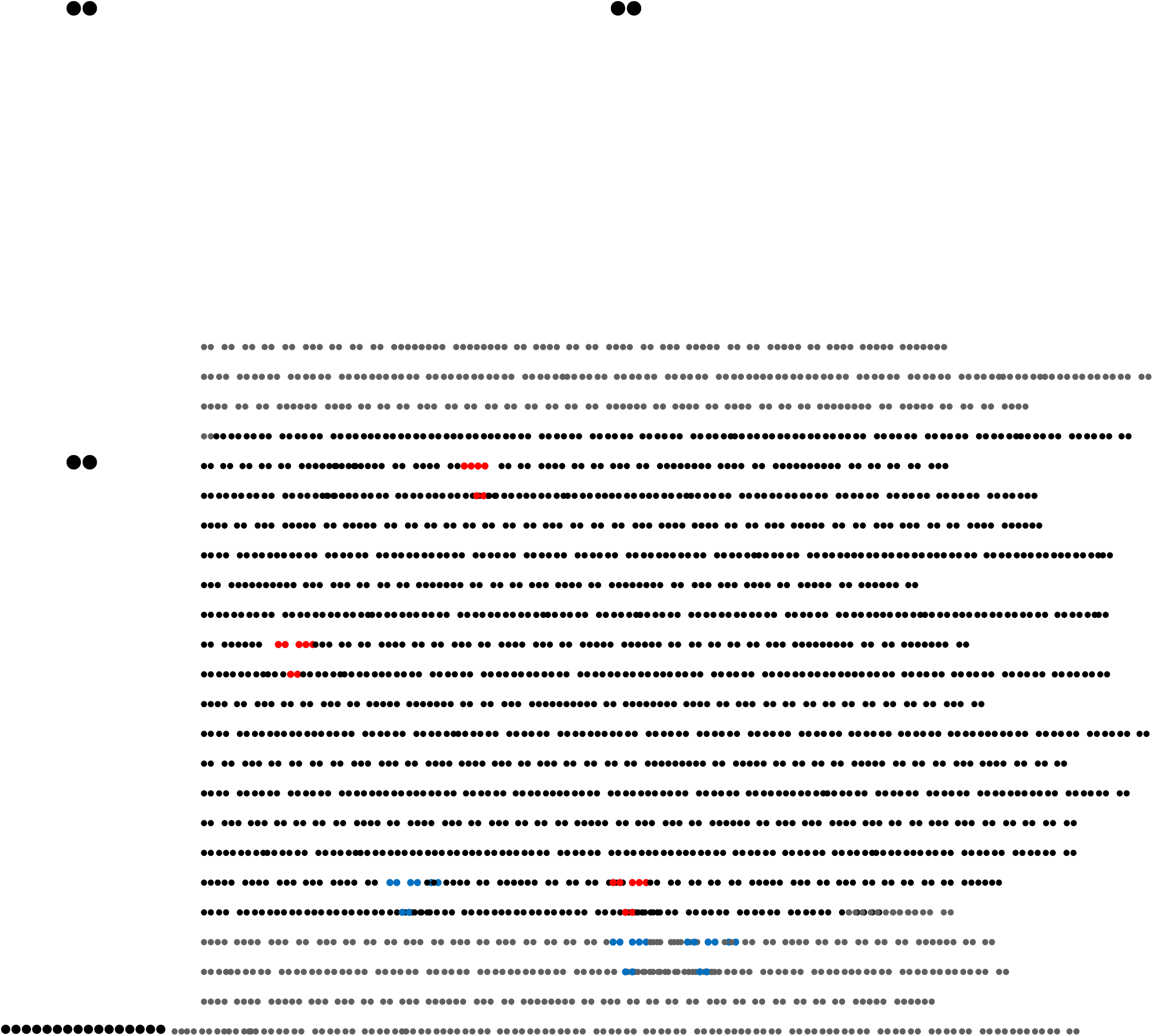
Representation of the 3D structure of TS 15-1c and prediction of active sites. The modeled structure of TS 15-1c is shown as a surface representation (A), and a Cα trace representation (B). The relative distribution of the surface charge is shown with acidic regions in red, basic regions in blue and neutral regions in white. Amino acid sequence of the C-terminal domain of TS 15-1 (C). Predicted active sites (H389, D444, and S533) and substrate binding sites (G527, S553, and G555) are shown in blue and red, respectively.

### Expression and localization of TS-15 in the *T. spiralis* life cycle

In order to determine when the *Ts 15-1* mRNA was the most highly expressed in the *T. spiralis* life cycle, real-time PCR was performed on new born larvae, adult worms, muscle stage larvae of *T. spiralis*, and during the *T. spiralis* infection period (at 0, 7, 14, 28 days after infection). As the results show, the *Ts 15-1* gene was the most highly expressed in muscle stage larvae, and its expression is also highly elevated 28 days after infection (Fig. 5A). In order to know whether TS 15-1 was secreted from parasites, an α-TS 15-1c antibody was produced and was reacted with *T. spiralis* ES-P and total extract. The TS 15-1c antibody strongly reacted with proteins around 72 kDa in ES-P and slightly reacted with a total extract at the same size (Fig. 5B). Furthermore, to know whether TS 15-1 has antigenicity or not, *T. spiralis* infected mice sera (0, 1, 2, and 4 weeks after infection) were reacted with recombinant TS 15-1c protein (Fig. 5C). rTS 15-1c most strongly reacted with mouse serum collected 4 weeks after infection. To know where TS 15-1 is secreted in the parasite, α-TS 15-1c antibody was reacted against serial sections of the *T. spiralis* muscle stage larvae using immunohistochemical methods. α- TS 15-1c antibody strongly reacted with the inner layer of the cuticle and the ladder shapes structure around the esophagus in muscle stage larvae that appear to be stichocytes (Fig. 6).

**Fig. 5.**
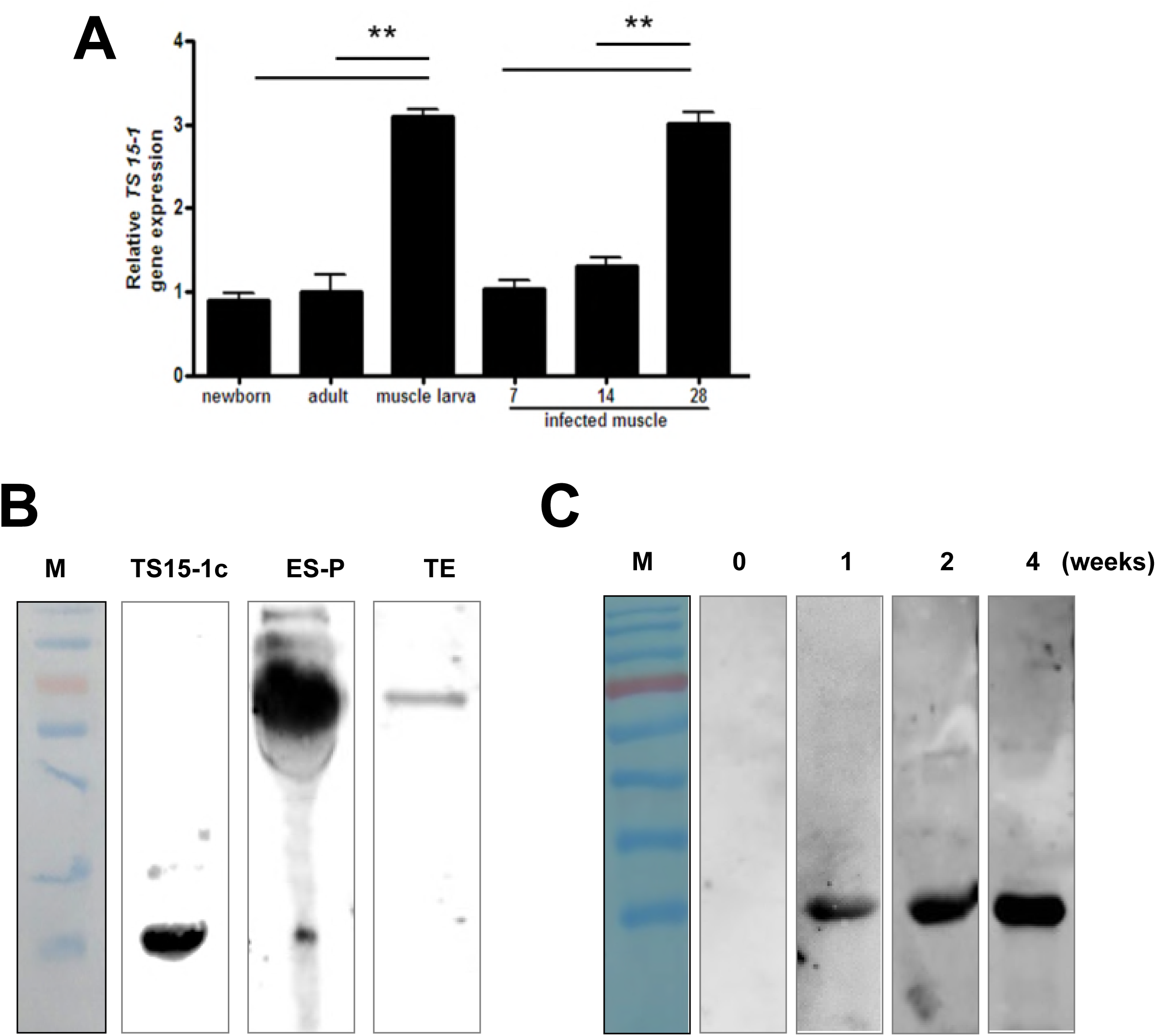
Expression levels of *TS15-1* during developmental stages and *T. spiralis* infected muscle, and evaluation of TS15-1 antigenicity. Larva from each stage, the adult worm, and *T. spiralis* infected muscle (1, 2 and 4 weeks after infection) were collected and *TS 15-1* gene expression levels were analyzed with real-time PCR analysis (A; **; P< 0.01, n = 5 mice/group, these were representative results from three independent experiments). Ten μg each of ES-P and total extract from *T. spiralis* were separated on SDS-PAGE and transferred to an immunoblotting membrane for western blot analysis. The polyclonal α-TS 15-1c antibody (1:500 dilution) was used as primary antibody (B; M, molecular marker). Ten μg of purified recombinant TS 15-1c protein was loaded on SDS-PAGE and transfer to an immunoblotting membrane for western blot analysis. The membrane reacted with the sera (1:500) of *T. spiralis* infected mice which were sacrificed at 1, 2 and 4 weeks after infection as primary antibodies (C; M, molecular marker).

**Fig. 6.**
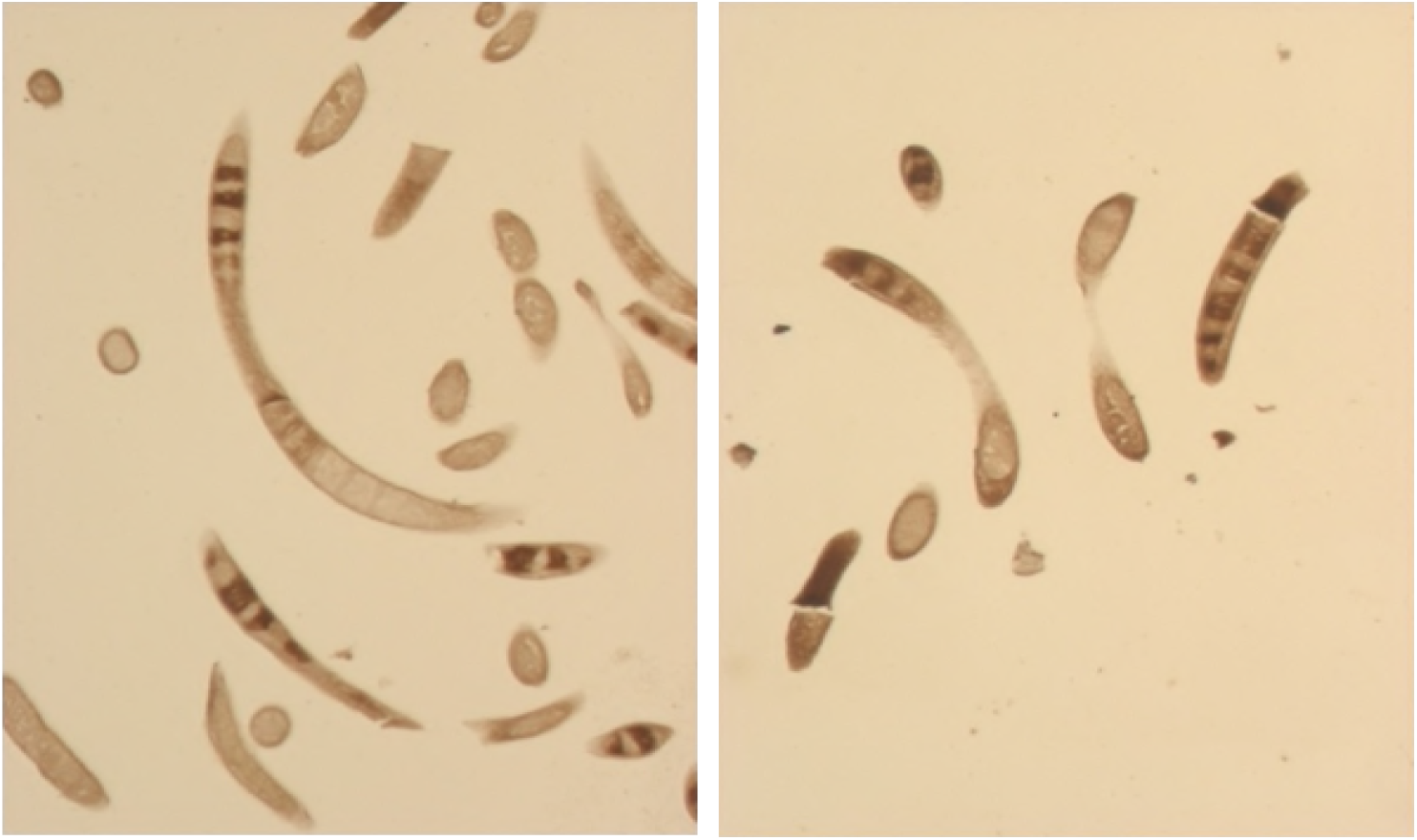
Localization of TS 15-1 in muscle stage larvae. Serial sections (5 - 6 μm) of muscle larvae embedded on paraffin blocks were reacted with α-TS 15-1c antibody.

### Evaluation of the TS 15-1c protein type I collagen inducing ability

In order to know whether TS 15-1c had type I collagen elevating ability, we applied TS 15-1c to the ear skin of 6 week-and 14 week-old mice and evaluated the expression levels of *collagen I*, *Smad2/3*, and *TGF-β1*. All of the tested genes’ expression levels, including the *collagen I* gene of the 14 week-old mice, were significantly lower than those in the 6 week-old mice. However, after 14 TS 15-1c treatments on 14 week-old mice, *collagen I, Smad2/3*, and *TGF-β1* gene expression levels in these mice were significantly increased compared with non-treated mice of the same age (Fig. 7A). We investigated protein levels of type I collagen, the phosphorylation form of Smad2/3, and levels of TGF-β1 in TS 15-1c treated 14 week-old mice and the protein levels were considerably recovered relative to those of the non-treated 14 week-old mice (Fig. 7B)

**Fig. 7.**
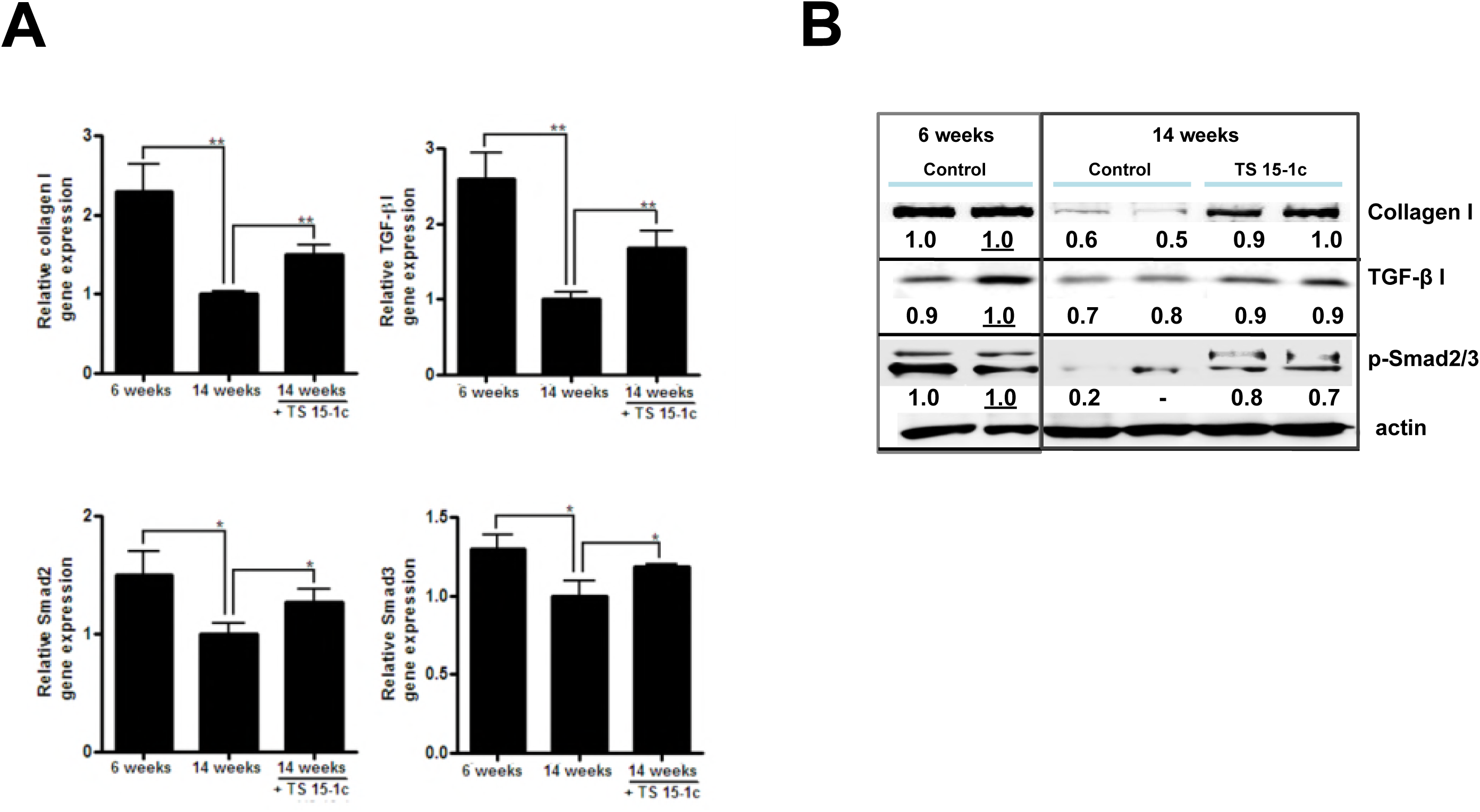
Evaluation of collagen I and TGF-β signaling pathway-related protein expression after treatment of recombinant TS 15-1c protein on natural aged mice. Schematic schedule of the purified TS 15-1c treatment (A). 14-week-old mouse ears were treated with purified TS 15- 1c protein (30 μg) daily for 14 days. On day 14, total RNA was extracted from each ear tissue and the gene expression levels of *collagen I*, *TGF-β1*, and *Smad2/3* were analyzed via real-time PCR. (B; *; *P* < 0.05, **; *P* < 0.01, n = 3 to 5 mice/group, these were representative results from three independent experiments). Total proteins were extracted from each mouse group for western blot analysis (C). 15 μg of each protein sample was loaded and separated in a 10% acrylamide SDS PAGE gel and transferred to nitrocellulose membrane. The α-TGF-β1, p-Smad2/3, α-mouse type I collagen and actin antibodies were incubated as primary antibody and α-mouse or α-rabbit IgG antibody conjugated with HRP was reacted as secondary antibody. After adding ECL substrate, the membranes were analyzed using the LAS 3000 machine. (Areas of the detected bands were determined and compared by Image J software).

## Discussion

In this study, we identified the host collagen inducing factor from *T. spiralis*, named it TS 15-1, and confirmed its serine protease activity and ability to elevate type I collagen, TGF-βI, and related signal proteins (Smad2/3) on a transcriptional and protein level. In addition, we found that it was expressed outside the parasite and elicited specific antibody production from the host immune system. In a previous study, we revealed that the ES-P of *T. spiralis* could induce collagen production of host muscle tissue during the infection period, and that it was closely related with serine protease activity 9].

We can carefully suggest that TS 15-1 is one of the key collagen inducing factors in ES-P revealed in our previous study. Although we could not demonstrate that TS 15-1 is one of the key molecules in the nurse cell formation step, it might be one of the central factors for nurse cell formation. This is because *TS 15-1* gene expression level was the highest during the *T. spiralis* muscle larva stage and its specific antibodies could be detected in mouse serum from 1 week up to 4 weeks after a *T. spiralis* infection (Fig. 5). During the nurse cell formation period (1 week - 4 weeks), *T. spiralis* might strongly secrete TS 15-1 to induce collagen capsule synthesis by the host muscle cell. In order to demonstrate TS 15-1’s collagen inducing ability, we treated naturally aged mice with TS 15-1c. Recombinant TS 15-1c protein could elevate collagen I production and the TGF-βI signaling pathway related to Smad2/3 proteins (Fig. 7). Type I collagen expression is closely related with the TGF-βI/Smad2/Smad3 signaling pathway [10-12].

Parasite secretory proteases might have important functions in modulating the interactions between parasites and hosts because of their particular roles in the invasion of host tissues, parasite nutrition, and evasion of host immune responses [13-16]. A trypsin-like serine protease of parasites could be involved in host immune regulation [17-19]. Serine proteases in nematodes are known to be involved in invasion into host cells and tissues and are likely to be important in molting [20]. TS 15-1 was revealed to be a serine protease, trypsin like protein, because its activity was inhibited by PMSF and it was composed two domains which were very similar but not identical to each other (Fig. 2C and Fig. 3A). Several secreted serine proteases have been identified among *T. spiralis* ES proteins, including the 69 kDa putative serine protease TsSerP (two trypsin-like domains), the 45 kDa serine protease TspSP-1, and a 35.5 kDa serine protease [13, 15, 21-23]. Most of these have strong antigenicity, specific antibodies for them are easily detected experimentally in infected animal sera, and they have one or two trypsin like domains [13, 15, 21]. Most secreted proteases could elevate their specific antibodies during nematodiasis [21, 24]. Trap et al., reported the identification of the putative serine protease, TsSerP, isolated from the *T. spiralis* adult-newborn larvae stage. It has two trypsin-like serine protease domains flanking a hydrophilic domain, which is the same structure as TS 15-1. Homology of the full length amino acid sequences of TsSerP and TS 15-1 was about 98% (data not shown). Immunohistochemistry analysis revealed that TsSerP was located on the inner layer of the cuticle and esophagus of the parasite, TS 15-1 was also detected on the inner layer of the cuticle and stichocytes in this study (Fig. 6). These two serine proteases of *T. spiralis* might have similar functions, although the function of TsSerP was not clearly revealed [13].

In this study, it was revealed that TS 15-1 could elevate collagen expression via the TGF-β1 signaling pathway in host tissue of normal aged mice. This characteristic could be used for therapeutic effects including wound healing and cosmetic usefulness with wrinkle reduction. The various serine proteases may participate in physiological or pathological processes, like tissue repair, vascular remodeling, and wound healing, that depend on cell proliferation and migration [25, 26]. Wound healing is a well-orchestrated process, where numerous factors are activated or inhibited in a sequence of steps [27]. Numerous signaling pathways are involved, among of them, the TGF-β1/Smad pathway is representative and well known to participate in the wound healing process [27]. Hozzein et al., suggested that topical application of propolis would promote the wound healing process by activation of TGF-β1 pathway [28]. The gradual loss of collagen in skin with aging results in wrinkles and other signs of skin aging [5]. The content of type I collagen, the major collagen in the skin and a marker of collagen synthesis, is deceased by 68% in old skin versus young skin, and cultured young fibroblasts synthesize more type I collagen than old cells [5].

In conclusion, we identified a host collagen inducing factor from ES-P using immune screening methods and demonstrated the molecular/genetic characteristics and function of TS 15-1. Further study will be required to understand the detailed mechanisms for receptors in the host cells, and to identify the minimal structure that can induce collagen for cosmetic and medical purposes.

---

**Supplementary Fig. 1.** Result of immunoscreening of *T. spiralis* cDNA library by α-F3 antibody. After *T. spiralis* cDNA containing phages were mixed with *E. coli*, the plaques were incubated with NC membranes for 4 hrs. The membranes were reacted with α-F3 antibody (1:500) as the primary antibody and α-rat IgG antibody conjugated with HRP was reacted as secondary antibody. After adding of 3,3'-diaminobenzidine (DAB), colored spots were compared with original plates. Positive plaques were amplified and re-analyzed by the same method.

